# Time-resolved structures of β_2_-adrenergic receptor modulation by a photoswitchable beta-blocker

**DOI:** 10.1101/2025.09.03.673938

**Authors:** Robin Stipp, Quentin Bertrand, Matilde Trabuco, Anna Duran-Corbera, Maria Tindara Ignazzitto, Hannah Glover, Fabienne Stierli, Juanlo Catena, Melissa Carrillo, Sina Hartmann, Hans-Peter Seidel, Matthias Mulder, Thomas Mason, Yasushi Kondo, Maximillian Wranik, Martin Appleby, Christoph Sager, Raymond Sierra, Gregory Gate, Pamela Schleissner, Xinxin Cheng, Tobias Weinert, Robert Cheng, Sandra Mous, John H. Beale, Michal Kepa, Amadeu Llebaria, Michael Hennig, Xavier Rovira, Joerg Standfuss

## Abstract

G protein-coupled receptors (GPCRs) regulate essential physiological responses and are important drug targets, yet their ligand-induced conformational dynamics remain poorly understood. The β_2_-adrenergic receptor (β_2_AR) is a prominent member of the GPCR family. It regulates bronchial and vascular function and is a significant drug target, particularly in respiratory and smooth muscle-related disorders. We employed time-resolved crystallography at X-ray free-electron lasers (XFELs) to capture the conformational dynamics of β_2_AR bound to photoazolol-1, a beta-blocker derivative developed for photopharmacological applications. Structural snapshots of the receptor bound to *trans*-photoazolol-1 (pre-photoconversion), a strained intermediate, and the fully photoisomerized *cis*-photoazolol-1 reveal an intricate interplay between ligand chemistry and receptor plasticity. Isomerization of the azobenzene moiety induces distinct conformational changes within the orthosteric pocket, altering interactions with the extracellular loop 2 and transmembrane helices 5 and 6. Supported by functional assays, these structural shifts suggest that photoazolol-1 transitions from an inverse agonist to a neutral antagonist upon photoactivation. Our findings uncover a mechanism of GPCR modulation reminiscent of rhodopsin activation and offer a framework for designing ligands that harness light-driven transitions to achieve spatiotemporal control of receptor function.

## Introduction

G protein-coupled receptors (GPCRs) are central to many human signaling pathways and are common targets in drug development^1^. In cases where disease disrupts natural homeostasis, many GPCRs serve as essential “adjustment knobs” to restore physiological equilibrium. Beta-adrenergic receptors are prominent GPCRs that regulate the body’s physiological responses to stress. Small-molecule drugs like propranolol, known as beta-blockers, inhibit adrenergic receptor activity by blocking the binding site for endogenous adrenaline and noradrenaline to reduce heart rate and blood pressure in patients with different cardiovascular issues. The specificity of these drugs for specific adrenergic receptor subtypes, along with their action as agonists or antagonists, is critical for minimizing off-target effects.

Numerous adrenergic receptor structures^2^ have been solved with the β_2_-adrenergic receptor (β_2_AR) acting as a reference to study GPCR activation^3^ due to the early availability of both ligand-bound^4-7^ and G protein-bound states^5^. Some of the secrets behind the adaptions of the protein to different ligands have been uncovered using multiple methods, including time-resolved cryo-electron microscopy^8^, nuclear magnetic resonance spectroscopy^9^ and molecular dynamics simulations^10^. Yet little experimental data shedding light on the whole structure at the atomic level during these adaptations is available, limiting our potential to understand receptor plasticity in presence of small molecule drugs and other ligands.

The ultrafast yet extremely bright pulses from X-ray Free Electron Laser (XFELs) sources enable the outrunning of structure-altering radiation damage and the resolution of conformational changes in proteins at physiologically relevant temperatures and with high spatiotemporal resolution. Serial femtosecond crystallography (SFX) using these ultrabright X-ray sources has been particularly successful in resolving GPCR structures from microcrystals directly grown in membrane mimicking phases^11,12^. Time-resolved femtosecond serial crystallography (TR-SFX)^13,14^ allows to resolve ultrafast structural snapshots at XFELs but has also been adapted to resolve slower intermediates at synchrotron sources^15,16^. The technique has been employed on a growing series of proteins^16,17^ and has been particularly successful with light-triggered proteins, including rhodopsin, the GPCR central to our visual system^18^.

Rhodopsin constitutes another important system to understand GPCR activation^19^ as it can be purified from native sources and can be triggered efficiently by an ultrafast photoreaction that alters the bound retinal chromophore from an inverse agonist to a full agonist. Photoactive ligands like retinal are ideal for time-resolved studies as they allow synchronized initiation of a specific process to be observed. However, they are not available for most GPCRs, prohibiting the direct study of ligand interaction dynamics on the structural level. Exceptions are created by the field of photopharmacology. The field focuses on chemically modifying ligands to put pharmacologically relevant targets under the control of light to achieve superior temporal and spatial control over target activity compared to conventional medications^20-22^. Due to their modifiable chemical scaffolds and high quantum yield^23,24^, azobenzene-based compounds have been the focus of many recent advancements, including several photoswitchable agonists^25^ and antagonists^26,27^ targeting adrenergic receptors.

One such photoswitch is photoazolol-1 (or PZL-1), a compound derived from propranolol, the first major β–receptor–blocking drug to be used clinically^28^. Similarly, to its parent compound, photoazolol-1 has a high affinity and acts as a potent beta-blocker based on cell-based cAMP reporter assays^27^. It retains the characteristic fingerprint region common to many synthetic β_2_AR ligands and the endogenous adrenaline and noradrenaline hormones but replaces the aromatic moiety, present in related β_2_AR antagonists, with a p-acetamido substituted azobenzene to allow for efficient switches between *trans* and *cis* configurations^27^. In its metastable *cis*-form, photoazolol-1 has been shown to have a considerably lower antagonistic potency than the *trans*-isomer^27^; however, whether this effect is solely based on a loss of affinity or a change in receptor-ligand interaction dynamics is not understood.

In this study, we determined the room-temperature structure of the photoazolol-1/β_2_AR complex at 2.5 Å resolution and investigated the effect of illuminating photoazolol-1 within its binding pocket using TR-SFX at the Linac Coherent Light Source (LCLS) and the Swiss X-ray Free Electron Laser (SwissFEL). Photoisomerization repositions the ligand within nanoseconds, but counter to our initial expectations, even after several seconds, photoazolol-1 does not leave the receptor. Instead, its p-acetamido azobenzene group finds a new position between transmembrane helix 5 and 6 (TM5 and TM6), where it disrupts the packing of the seven transmembrane helical bundle and induces adaptive conformational changes in nearby residues related to receptor activation. Cellular assays, together with comparisons of these rearrangements to early changes in the photoactivation of visual rhodopsin, suggest that *trans-* photoazolol-1 has inverse agonist activity that is abolished upon illumination, offering new insights into the molecular basis of photopharmacology and receptor activation.

## Results

### Photoazolol-1 binding mode

One key challenge in photopharmacology is designing photoswitchable ligands that retain the binding mode of the parent compounds in one of their configurations. To compare the binding mode of *trans*-photoazolol-1 to conventional beta-blockers, we first solved the crystal structure of β_2_AR bound to dark-equilibrated photoazolol-1 (**Figure 1A**). Notably, even after thorough optimization our crystals diffracted only to low resolution using conventional cryo-crystallography at synchrotrons, as is not uncommon for GPCR microcrystals^12^. High-intensity XFEL pulses were essential to reach resolutions of 2.5 Å (**Supplementary Table 1**), which are typically required to accurately model ligand and water-mediated interactions for structure-guided drug design applications. The inability to grow large, well-diffracting crystals may be due to the chosen construct that contains a T4 lysozyme fusion and terminal truncations to facilitate crystallization but only a single mutation (compare material and methods), as systematic thermostabilization approaches often have the drawback to limit conformational flexibility of the targeted receptor^29^. A root-mean-square deviation of Cα atoms of 0.638Å to the propranolol-bound β_2_AR structure from LCLS^30^ shows the photoazolol-1-bound structure is near identical, confirming the successful design of the photoswitchable compound. The high degree of similarity (**Figure 1B and 1C)** includes the fingerprint region of photoazolol-1, which interacts with the receptor binding pocket through the same interactions as propranolol, including the characteristic polar contacts with Asp113^3.32^ and Asn312^7.38^ [superscripts denote Ballesteros-Weinstein numbers^31^], similarly found in other adrenoceptor-ligand complexes^32,33^ and known to be critical for potency and efficacy of agonist activation^34,35^.

**Figure 1:**
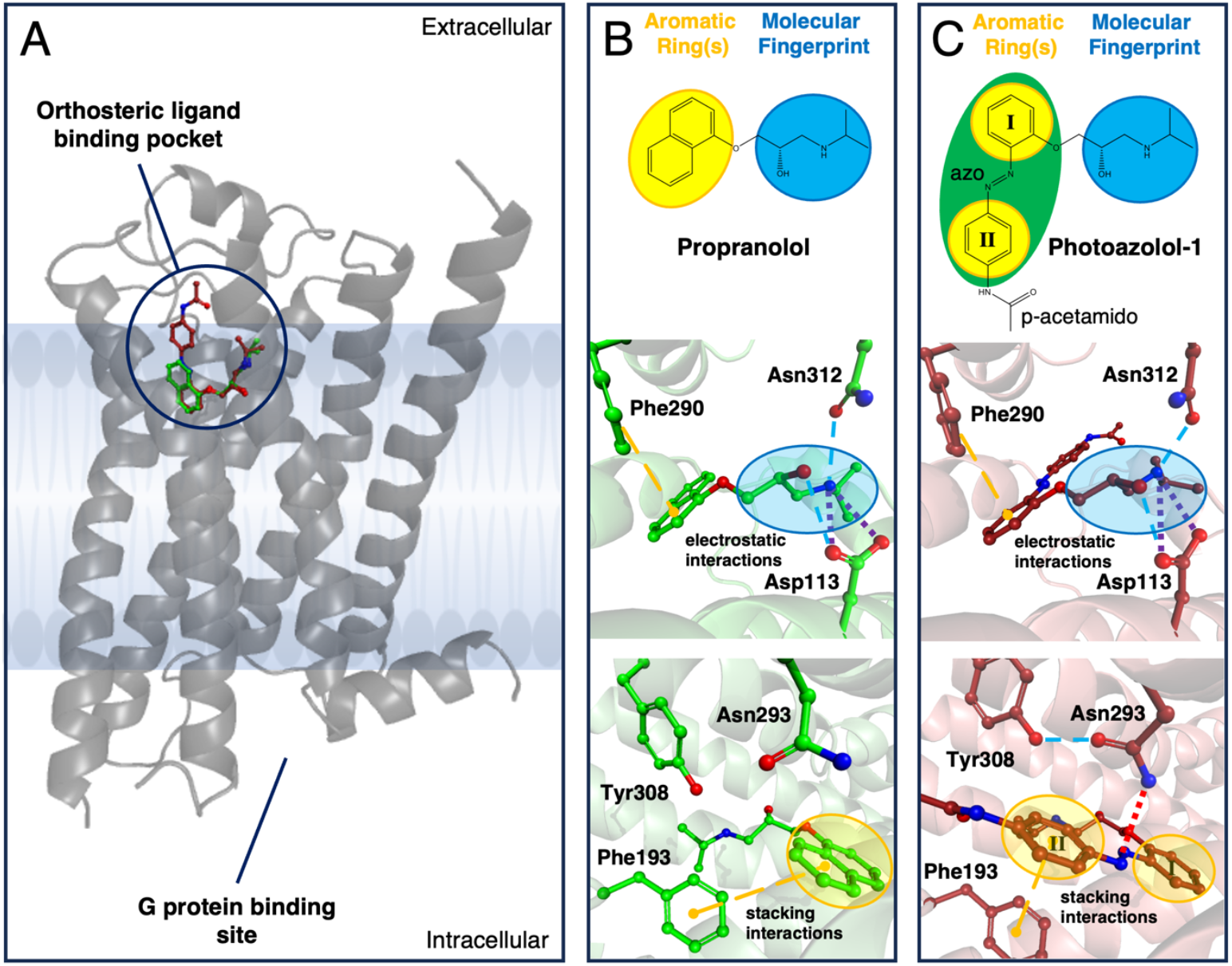
Comparison of propranolol and photoazolol-1 binding. (**A**) The β_2_AR follows the prototypical 7TM fold (grey cartoon) of GPCRs with the orthosteric ligand binding site (Propranolol (green) and Photoazolol-1 (red) are shown in ball and sticks representation) close to the extracellular and the G protein-binding site on the intracellular side of the membrane. (**B**) The beta-blocker propranolol (6ps5)^30^ contains a molecular fingerprint region (blue) and aromatic rings (yellow) that bind the β_2_AR with high affinity through a series of ionic (dotted lines) and hydrophobic stacking (dashed lines) interactions. (**C**) Photoazolol-1 maintains the general chemical structure of propranolol but separates the two rings in a photoswitchable azobenzene group (green) with an acetamido group on the para position of ring II. The main interactions of the original beta-blocker with the β_2_AR binding pocket are maintained despite the modifications in the synthetic photoswitch.

Interestingly, the photoswitchable azobenzene moiety maintains π–π interactions with Phe193^45.52^ and Phe290^6.52^, despite a doubling in the distance between the aromatic rings in photoazolol-1 compared to propranolol. A difference in the binding mode of *trans-*photoazolol-1 compared with propranolol, is the additional polar interaction between the azobenzene bond of the former molecule and Asn293^6.55^, which changes the rotamer to establish this contact. This interaction is further stabilized by an additional hydrogen bond between Asn293^6.55^ and Tyr308^7.35^. Interestingly, this tyrosine is replaced by a phenylalanine in the β_1_-adrenergic receptor, suggesting a degree of subtype specificity as observed for the related opto-prop-1 molecule^26^. The overall similar binding mode of *trans*-photoazolol-1 is reflected in a comparable binding affinity: 9.8 nM for propranolol and 14.1 nM for dark-adapted photoazolol-1 (**Supplementary Figure 1**). The design of photoazolol-1 thus resulted in a compound where both binding mode and binding affinity are similar to other known beta-blockers, fulfilling one of the key requirements for an efficient photochemical trigger of β_2_AR activity.

### Effect of photoazolol-1 illumination

Another key requirement for an effective photopharmacological compound is its ability to induce a change in its biological effect upon illumination. For photoazolol-1, irradiation with 380 nm light results in a 17-fold reduction in antagonistic activity, even though only 86% of the compound converts to *cis*-photoazolol-1 in the photostationary state^27^. To investigate the molecular basis of this effect, we performed time-resolved serial crystallography experiments during two beamtimes at the Linac Coherent Light Source (LCLS) and the Swiss X-ray Free Electron Laser (SwissFEL). At LCLS, we employed an injector-based approach to probe an early time delay of 17 ns, twice the pulse length of the used optical laser. Based on previous experience with retinal proteins^18,36,37^ and the ultrafast kinetics of azobenzene isomerization^38,39^, we expected to resolve the initial effect of photoazolol-1 isomerization on the receptor. Such a classical TR-SFX approach is, however, unsuitable for longer time delays as it is limited by extrusion stability and the achievable distance between pump-probe pulses. At SwissFEL, we therefore relied on a fixed-target approach to resolve the effects on the receptor approximately 10 s after photoactivation. The differences and similarities between the two experiments are further explained in **Supplementary Figure 2**. Progressing from the starting point of the *trans*-photoazolol-1 structure, isomorphous difference electron density maps [*F*_o_(light) − *F*_o_(dark)] enabled us to track changes in protein-ligand interactions (**Supplementary Figure 3**) over time. To further analyze these structural changes, we determined extrapolated structure factors using an activation level of 28% for the 17 ns and 15% for the 10 s time delays. While we cannot exclude an ensemble of structures with lower occupancies, single molecular structures refined well against the extrapolated data. The potential impact of data extrapolation could be further limited by combining the dark and light models and refining them against the overall data at each time delay. A morph between the structure highlights the conformational changes in the binding pocket and along 7TM helical bundle (**Supplementary Movie 1**).

The structures show that photoazolol-1 adopts a strained *cis* configuration 17 ns after illumination (**Supplementary Figure 4**), leading to only minimal changes in the receptor (**Figure 2**). Unable to chemically relax within the tight constraints of the binding pocket, the photoswitch maintains much of its original shape. The light-induced isomerization, however, causes a shift in the p-acetamido azobenzene moiety by 4.2 Å (measured between the respective positions of the CO2 atom), apparently weakening interactions with the receptor. Specifically, the π–π interactions with Phe290^6.52^ are lost, and increased distances with less favorable geometry suggest weakened interactions with Phe193^45.52^ and Asn293^6.55^. Despite these alterations, the ligand remains anchored through strong polar interactions mediated by the fingerprint region. Given the size limitations of the binding pocket and the minimal changes in the receptor during this early time delay, it seems likely that the initial photoreaction proceeds without a substantial rotational component around the azo bond. These early rearrangements leave the ligand in a metastable pose with its stored strain energy preserved to induce conformational changes in the protein – providing an observation window into the dynamic response of a GPCR to its bound ligand – similar to the response of rhodopsin after initial photoisomerization^18^.

**Figure 2:**
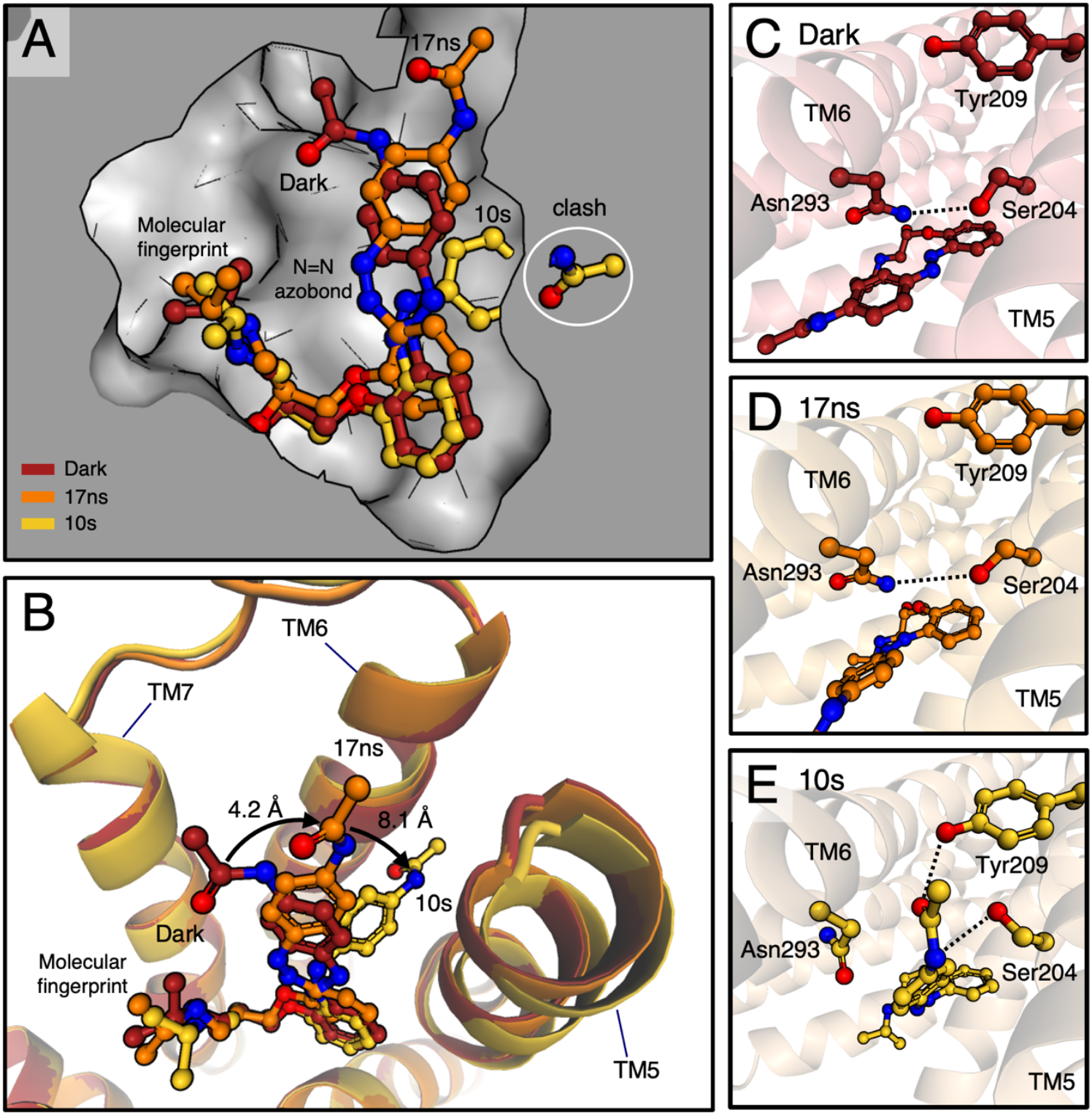
Structural effects of photoazolol-1 isomerization. (**A**) The binding pocket of the β_2_AR dark state (grey) with superposed photoazolol-1 (dark state in red, 17 ns in orange and 10 s in gold). The initially twisted *cis*-photoazolol-1 fits the binding pocket, while relaxation at the later time delay would lead to a clash with the interface between TM5 and TM6. (**B**) Photoazolol-1 isomerization leads to a relocation (black arrow) of the p-acetamido substitutions into a new position between TM5 and TM6. Superposition of the dark (red) and 10 s structures (gold) show how the binding pocket adapts through an outward move of TM5 and TM6 combined with an inward shift of TM7. Oxygen atoms are shown in red and nitrogen atoms in blue. In the dark (**C**) and **after** 17 ns (**D**), the ionic interactions (dotted lines) between Ser204^5.43^ and Asn293^6.55^ are maintained, while full photoazolol-1 isomerization after about 10 s (**E**) interrupts this interaction implicated in agonist-induced β2AR activation.

In our previous studies investigating protein-ligand interaction dynamics in tubulin^40,41^, the adenosine A2a receptor^42^ and the metabotropic glutamate receptor 5^43^, the synthetic photoswitches left the binding pocket within milliseconds after activation. In contrast, photoazolol-1 remains bound to the β_2_AR even after 10 s, three orders of magnitude longer than we could probe in our previous experiments. However, within 10 s after illumination the initially strained *cis*-photoazolol-1 configuration relaxed into a more favorable geometry (**Figure 2**) with slightly higher total energy than the original *trans*-state (**Supplementary Figure 4**). These energetic considerations are consistent with ligand retention in the binding pocket several seconds after isomerization. After relaxation of the transient photoazolol-1 configuration, the p-acetamido group, measured between the two CO2 labelled atoms, has shifted further with a maximum distance of 8.1 Å. The group finds a new position between TM5 and TM6, where it leads to a rotamer change of Ser204^5.43^ and Asn293^6.55^, interrupting the interaction between these two residues and establishing new interactions with Tyr209^5.48^ and Ser204^5.43^. The repositioned ligand has shifted the ends of TM5 and TM6 away from the original binding pocket, whereas TM7 moves in on the other side, keeping the overall volume of the binding pocket constant. Comparisons between structures containing ligands with various pharmacological profiles have implicated similar interactions with residues in TM5 and TM6 in the agonist-induced activation of beta-adrenergic receptors^32,44-46^. However, despite the local rearrangements in the binding pocket, general structural characteristics of GPCR activation (including the DR^3.50^Y, CW^6.48^xP, and NP^7.50^xxY motifs) remain in an inactive conformation. As in other structures without bound signaling partner, the local changes upon photoazolol-1 isomerization did not propagate into larger changes on the intracellular side where the G protein binds.

### Molecular mechanism of photoazolol-1 activity

A guiding principle of azobenzene-based approaches to photopharmacology, is the design of small molecules with differential function between *cis* and *trans* configurations. Our time-resolved observations shed light on this mechanism with photoazolol-1, a ligand that remains in the binding pocket but exerts striking differential activity upon photoisomerization. To investigate this observation, we characterized its dark and light-activated forms in the cellular context (**Figure 3A**). Our results reveal that photoazolol-1 binding to the β_2_AR is stable, showing no evidence of unbinding under any of the tested conditions, confirming what was observed in the structures. Indeed, competitive assays suggest that photoazolol-1 remains bound to the receptor for over an hour following a thorough washing protocol, regardless of whether the samples were kept in the dark or illuminated (**Figures 3B, 3C and Supplementary Figure 1A**). These results are very similar to those found for propranolol (**Supplementary Figure 1B**) and the previously described opto-prop-1, a chemically related molecule without the p-acetamido group, which did not undergo significant UV-induced unbinding either. In contrast, opto-prop-2, the meta-substituted photoswitchable beta-blocker, demonstrated light-dependent unbinding^26^. Although photoazolol-1 remains in the receptor pocket, a right shift of the dose-response curve is observed in functional assays when cells are irradiated with a 380 nm light, showing a significant 53-fold change in potency (**Figures 3D, 3E and Table 1**). This could indicate that the isomerization to the *cis* isomer results in a loss of photoazolol-1 antagonist activity, even though the ligand remains bound. Of note, no change in potency is observed for the control antagonist propranolol upon illumination (**Supplementary Figure 5 and Table 1**). Strikingly, this functional antagonism is restored one-hour post-illumination, without additional ligand application (**Figure 3E**), suggesting reversibility through thermal relaxation back to the *trans* isomer. Very similar results were obtained when competing with higher concentrations of cimaterol (1 µM) for both photoazolol-1 and propranolol (**Supplementary Figure 6**). Of note here is that previous studies demonstrated that photoazolol-1 relaxes to the *trans* isomer more rapidly in functional assays on β2AR-expressing cells than in aqueous solution *in vitro* [Figure S11 in Duran-Corbera et al. 2020^27^], thus supporting the hypothesis that it does not leave the binding site and that the *cis* form is less stable than the *trans* in the context of the receptor.

**Table 1:** Pharmacological data. PPS refers to Photoinduced Potency Shift, which is the relation (fold shift) between the measured half maximal effective concentration (expressed as negative logarithm pEC_50_) in light and dark conditions. Standard errors of the mean (SEM) from three independent experiments measured in duplicate are given. Statistical differences from EC_50_ values between experiments measured right after and 1 hour after illumination are denoted for adjusted p-values as follows: ****p < 0.0001.

| Compound | DARK |  | LIGHT |  | PPS<sup>[a]</sup> | SEM |
| --- | --- | --- | --- | --- | --- | --- |
|  | pEC<sub>50</sub> | SEM | pEC<sub>50</sub> | SEM |  |  |
| Photoazolo-1 | 7,73 | 0,03 | 6,01<sup>****</sup> | 0,04 | 52,81 | 4,16 |
| Photoazolo-1 (+ 1 hour) | 7,51 | 0,02 | 7,31 | 0,09 | 1,65 | 0,14 |
| Propranolol | 8,08 | 0,09 | 7,76 | 0,07 | 2,08 | 0,08 |
| Propranolol (+ 1 hour) | 8,02 | 0,03 | 7,85 | 0,05 | 1,50 | 0,11 |

**Figure 3:**
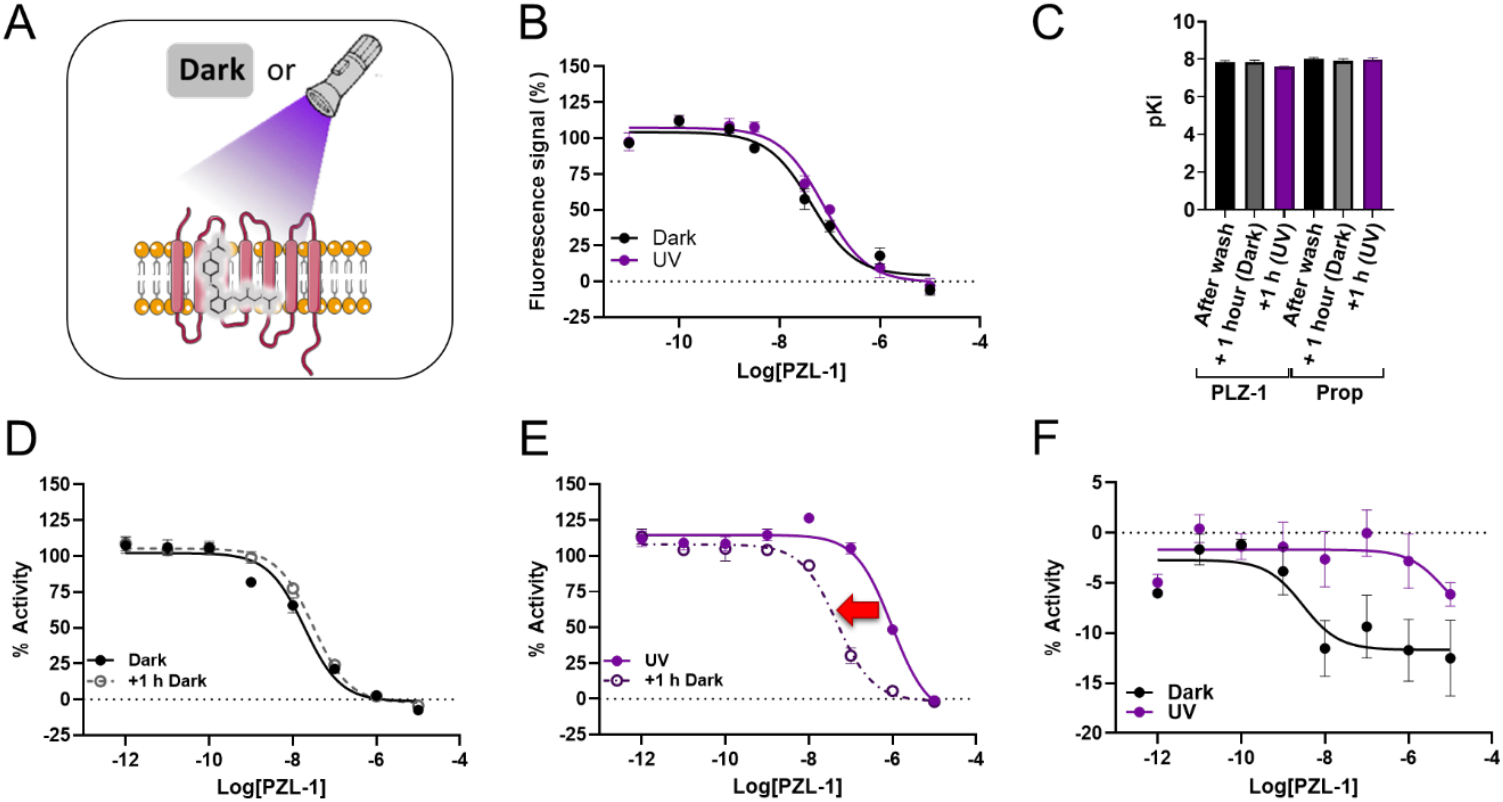
Light-dependent binding and activity of photoazolol-1 (PZL-1) on the β_2_AR. **(A**) Schematic representation of the pharmacological protocol in which cell cultures were either kept in the dark or exposed to light for 15 minutes following a 1-hour incubation with ligands. A thorough wash was then performed to remove any residual ligand from the medium. **(B)** Competitive binding curves of photoazolol-1 with a constant concentration of the fluorescent ligand carazolol-KK114 (100 nM). Measurements were performed 1 hour after washing. **(C)** Plot of the negative logarithm of the inhibition constant (pKI) values as calculated from the competitive binding curves. **(D)** Dose-response curves of photoazolol-1 with a constant concentration of the agonist cimaterol (10 nM) 15 min after washing kept in the dark (solid black lines), and after an additional 1-hour incubation in the dark (dashed black lines). **(E)** Dose-response curves of photoazolol-1 with a constant concentration of the agonist cimaterol (10 nM) 15 min after washing kept under light at 380 nm (solid violet lines) and after an additional 1-hour incubation in the dark (dashed violet lines). **(F)** Dose-response curves of photoazolol-1 in transfected cells to overexpress β2AR in the absence of agonist and 15 min after washing in the dark or under illumination with light at 380 nm. Data are shown as the mean ± standard error of the mean (SEM) of three independent experiments in duplicate.

To demonstrate that photoazolol-1 is a light-sensitive antagonist, we performed functional experiments in the absence of the agonist cimaterol. In these assays, no agonist activity of photoazolol-1 was observed when applied to cells overexpressing the β_2_AR (**Figure 3F**). On the contrary, photoazolol-1 decreased the basal activity, thus acting as inverse agonist in the dark. This effect was abolished upon illumination, indicating that the *cis* isomer acts as a neutral antagonist instead. This difference in functional activity between the *trans* and *cis* isomers is consistent with our structures, showing that the *cis* isomer adopts a distinct binding pose, altering key interactions within the binding pocket. Notably, the establishment of new contacts with Ser204^5.43^ and Tyr209^5.48^, and the loss of contact with Phe193^45.52^, a residue implicated in receptor activation and ligand efficacy, correlates with this differential activity (**Figures 1C and 3E**). Indeed, Phe193^45.52^ and adjacent residues have been highlighted in multiple studies as critical positions for ligand specificity and activity modulation [*e*.*g*., constrained catecholamines ^47^; β_1_ vs. β_2_ selectivity^48^; salmeterol binding^46^]. A recent study based on azo variants of clenbuterol has reported a photoinduced efficacy shift from partial agonist (*trans*) to antagonist (*cis*)^49^, providing further evidence that β_2_AR activity can be modulated by changing interactions within the binding pocket in response to light.

Possible explanations to reconcile the persistence of photoazolol-1 in the binding pocket with the light-dependent loss and recovery of functional activity could be the following. It is plausible that photoazolol-1 remains in the orthosteric pocket while exposing or allowing access to a secondary binding site for agonists only when bound in its *cis* state. This could be due to the loss of interaction between *cis*-photoazolol-1 and Phe193^45.52^, a residue potentially involved in the pocket reorganization and the agonist coordination at a secondary site within the entrance vestibule located between the extracellular loop 2 (ECL2), TM5 and TM6. Importantly, many changes are observed in the residues located in this region between the structures in complex with the *trans-* and *cis*-photoazolol-1. These include amino acid rotamer differences of His93^2.64^, His296^6.58^, Lys305^7.32^, Tyr308^7.35^ and Asp192^45.51^. This secondary binding site has been described for norepinephrine^50^. In addition, the exosite is occupied by the aryloxyalkyl tail of salmeterol as observed in the β_2_AR co-crystalized structure^46^. Interestingly, this site is also engaged by homobivalent bitopic ligands composed of two alprenolol-based β-blockers connected via a long linker^51^. Moreover, similar mechanisms have been described in other GPCRs for allosteric molecules binding at this location, such as muscarinic acetylcholine receptors^52^. Therefore, structural, binding and functional evidence support the hypothesis of the agonist binding into a second site at the same time as *cis*-photoazolol-1, a combination that would induce the activation of the receptor. An alternative explanation is that photoazolol-1 temporarily dissociates from the receptor, but remains in a non-washable, membrane-adjacent compartment, from which it can rebind. This mechanism has been described for other ligands targeting the β_2_AR^53,54^. Finally, we cannot rule out the possibility of the observed functional changes being driven by *cis*-specific allosteric effects from interacting proteins or from the existence of distinct receptor populations that differentially respond to specific ligand isomers. However, these two last-mentioned hypotheses would not fit with the time-resolved crystallography structural data, which suggest a stable binding of photoazolol-1 in both states within the orthosteric site of β_2_AR.

### Comparison to visual rhodopsin

In order to understand the effect of photoazolol-1 isomerization on activity, we compared the β_2_AR to visual rhodopsin, the GPCR activated through isomerization of its native ligand retinal within the binding pocket. Investigating the parallels could provide insights into the structural plasticity of GPCRs when reacting to a sudden change of a bound ligand and the mechanisms underlying the properties displayed by photoazolol-1 inactivation (**Figure 4**). The overall structural changes after light activation can be compared by overlaying the structures of dark^18^, lumi^55^, and activated meta-II rhodopsin^56,57^ with those of our time-resolved β_2_AR snapshots. Reminiscent of what we observed for photoazolol-1, the light-induced isomerization of the retinal polyene chain in rhodopsin places the β-ionone ring between TM5 and TM6. Ultimately, this intercalation results in a shift of 2.6 Å in TM6 and a smaller 1.6 Å shift in TM5, while TM7 moves inwards by 1.2 Å, changes very comparable to our structure containing the fully isomerized photoazolol-1 (**Figure 4A and 4B**). A further analogy in both receptors is the anchoring of the ligand to the protein at the part of the molecule most distant from TM5 and TM6. While this is achieved in the β_2_AR *via* the conserved interactions of Asp113^3.32^ in TM3 and Asn312^7.38^ in TM7 to the fingerprint region, the same is achieved in rhodopsin through electrostatic interactions to the retinal counterion residue Glu113^3.28^ in TM3 and the covalent Schiffbase to Lys296^7.42^ in TM7. While these overall changes within the binding pockets are strikingly similar, there are also significant differences, including larger changes in TM3 in rhodopsin and the outward-directed movement of the cytosolic ends of TM5 and TM6 that is characteristic for the G protein-binding conformation. Further conformational changes may have been inhibited by crystal packing but most GPCRs, including the β_2_AR, need both an agonist and a signaling partner to adopt a fully active conformation^10,58,59^.

**Figure 4:**
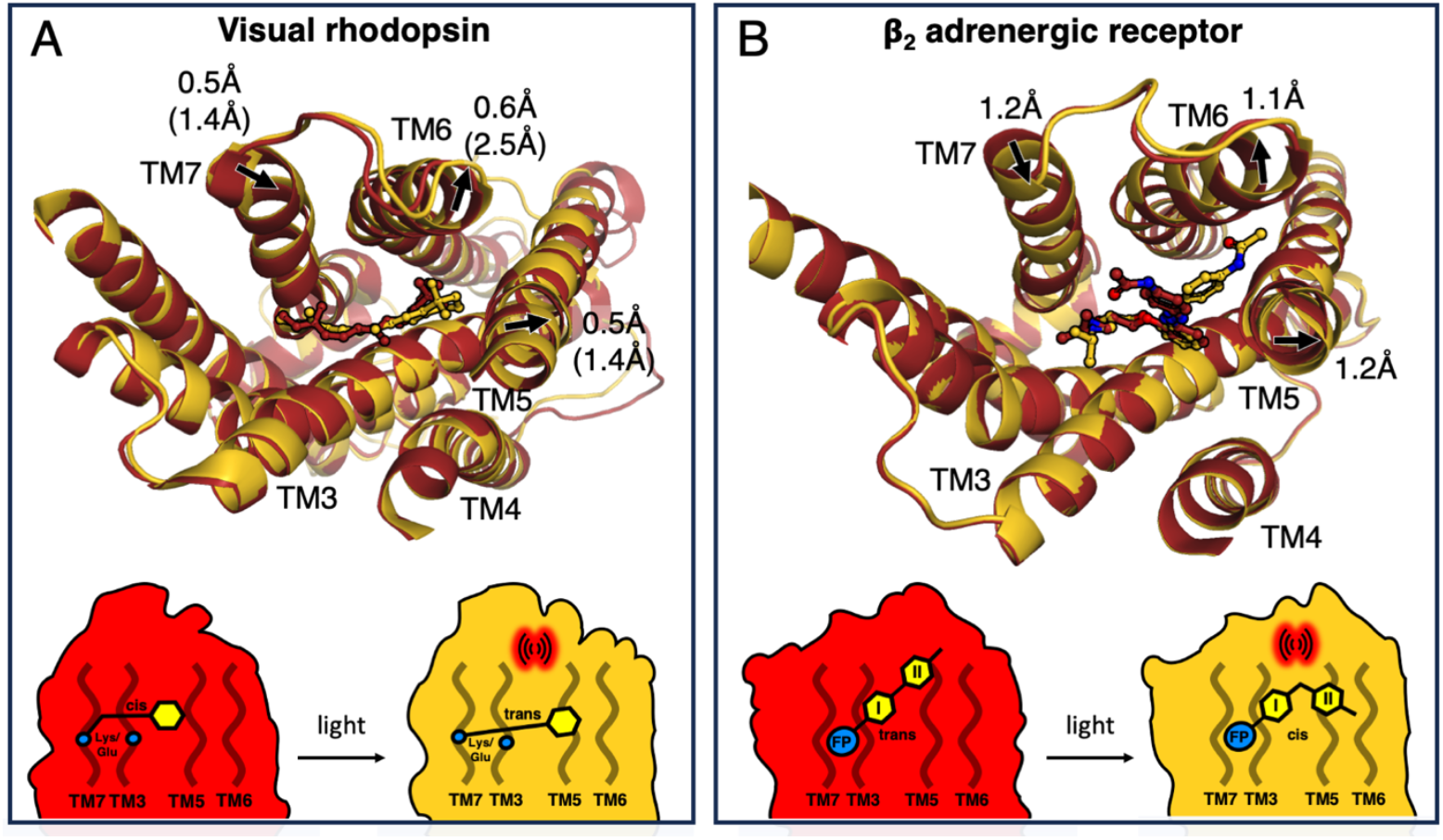
Comparison between rhodopsin and the β_2_AR. (**A**) The overlay of rhodopsin structures in the dark (pdb code: 7zbe) and lumi states (pdb code: 2hpy) shows how *cis*-to-*trans* isomerization relocates the β-ionone ring of retinal. This leads to shifts in the positions of TM5, TM6, and TM7, which are the initial steps towards reaching the fully activated metarhodopsin-II state (corresponding helix shifts in brackets). The schematic drawing shows the principal activation mechanisms with retinal covalently bound to Lys296^7.42^ in TM7 and interacting with TM3 by the Glu113^3.28^ counterion (blue circles). Light activation disrupts the local packing of the binding pocket through a shift of the β-ionone ring (yellow hexagon) towards TM5 and TM6. (**B**) The overlay of the β_2_AR structures with photoazolol-1 in the *trans* and *cis* configurations indicates similar shifts in TM5, TM6 and TM7. The schematic drawing highlights the similarities to rhodopsin with the fingerprint region (blue circle) bound between TM3 and TM7 and *trans*-to-*cis* isomerization shifting ring II of including the p-acetamido group into a new position between TM5 and TM6, thereby interrupting the original packing of the binding pocket.

Typically, GPCR agonists bind more tightly to the active state than the inactive state^60^, and agonist binding is often considered to occur by conformational selection^19^. Unliganded opsin expresses the same behavior, where constitutively active variants can be directly activated by all-*trans*-retinal^61,62^. However, native rhodopsin is exceptional among GPCRs as it functions through an induced fit mechanism, where the prebound *cis* retinal acts as an inverse agonist that is transformed by light to induce activation. This activation mechanism is surprisingly flexible, as the Lys residue covalently linking retinal can be moved to other locations while maintaining the ability to activate the G protein in a light-dependent manner^63^. Covalently bound agonists have previously been used to study the mechanisms of β_2_AR activation^58,64^ and given the overall similarities in how Class A GPCRs are activated^35,65^, it seems feasible that disrupting the binding pocket and repositioning the ligand can modulate β_2_AR activation.

## Discussion

GPCRs function is now recognized to involve complex conformational dynamics that extend far beyond classical two-state activation models. Within this framework protein–ligand interaction dynamics play a central role in modulating receptor activity and the pharmacological potency of small-molecule drugs. A key challenge in developing beta-blockers has been the design of ligands that selectively antagonize adrenergic receptors while minimizing off-target or intrinsic pharmacological effects. Photopharmacology introduces light as a noninvasive tool for precise spatiotemporal control over drug activity. This is typically accomplished through photochemical affinity switches that undergo light-dependent changes in binding affinity, enabling reversible control of receptor function *via* dynamic ligand binding and release.

Our structural findings suggest that the photoswitchable beta-blocker photoazolol-1 may modulate the β_2_AR through a distinct mechanism. Rather than dissociating upon photoisomerization, the ligand remains bound and induces local rearrangements within the orthosteric binding pocket. This conformational perturbation favors receptor activation, representing a conceptual shift from reversible occupancy-based modulation (i.e. a photochemical affinity switch) to internal structural repositioning of protein-ligand interactions (i.e. a photochemical efficacy switch). Such an approach to modulate receptor activity by light offers several advantages, including avoiding intramolecular competition between *cis* and *trans* isomers and reduced ligand loss due to diffusion or dilution, ultimately resulting in a more sustained and efficient light-induced response.

These findings underscore the potential of photopharmacological ligands as mechanistic probes for understanding GPCR activation. When combined with time-resolved structural approaches, this strategy provides high-resolution mapping of the conformational landscapes that underlie ligand-induced modulation of receptor activity. Iterative structural studies using chemically diverse, light-responsive ligands may further reveal the intrinsic plasticity of GPCR binding pockets, informing the design of next-generation therapeutics.

## Materials and Methods

### Construct and β_2_AR expression

A previously described construct was used to obtain a version of the β_2_-adrenergic receptor suitable for crystallization and ensure suitable protein expression and stability^30,66,67^. Compared to the human wtβ_2_AR, the following modifications were performed on the construct (UniProt ID P07550): The C-terminus was truncated at residue 384 and ICL3 (residues 231-262) was replaced with a cysteine-free T4-lysozyme, improving stability and crystallizability. Additionally, a point mutation (E122W)^68^ was inserted to improve stability and protein yield. The construct was expressed in *Trichoplusia ni* Hi5 cells using the FlashBAC system. The cells were grown in Sf900 II media at a density of 2*10^6^ cells/ml at 27°C under constant shaking. Following Bacmid infection using 1% VOI, the cells were incubated for 72h before harvesting at 3500×g for 10min and flash freezing in liquid nitrogen.

### Protein purification

Membrane preparation consisted of resuspension of the thawed xcell pellet in hypotonic buffer (10 mM HEPES pH 7.5, 20 mM KCL, 10 mM MgCl_2_, supplemented with Pierce^TM^ Protease inhibitor tablets) in a 1:6 ratio of pellet mass to buffer volume. The cell membrane was cracked using a Dounce homogenizer (20 strokes) followed by ultracentrifugation at 200,000x g for 30 min. The supernatant was discarded, and the pellet was resuspended in fresh hypotonic buffer using an Ultra-Turrax T25 basic (10 seconds of mixing), followed by ultracentrifugation. This washing method was repeated twice using hypertonic buffer (10 mM HEPES pH 7.5, 20 mM KCl, 10 mM MgCl_2_, 1 M NaCl, supplemented with protease inhibitor). After the last centrifugation step, the pellet was flash-frozen and stored at -80°C.

The purification method is based on previous work^69^ but was modified for increased efficiency and purity. The membrane pellet was thawed on ice and resuspended in solubilization buffer (25 mM HEPES pH 7.5, 300 mM NaCl, 5 mM imidazole pH 7.5). To the solution, 2 mg/mL iodoacetamide, protease inhibitor tablets and DNase I was added. Lauryl Maltose neopentyl glycol (LMNG) and cholesteryl hemisuccinate (CHS) were introduced to a final concentration of 1% and 0.2% (w/v), respectively. The solution was stirred at 300 rpm and 4°C for 90 min and subsequently clarified by ultracentrifugation (200,000 g, 40 min). The supernatant was combined with TALON-beads by Takara Bio (previously equilibrated to the solubilization buffer) and incubated at 100 rpm and 4°C, stirring for 1 hr. 1 mL of TALON beads were used for every 100 mL of solubilization supernatant.

After incubation, the beads were packed into an XK16 column and washed with 10 column volumes (CV) of wash buffer I [50 mM HEPES pH 7.5, 300 mM NaCl, 20 mM imidazole pH 7.5, 0.03% (w/v) LMNG, 0.003% (w/v) CHS], followed by 5 CV of wash buffer II [50 mM HEPES pH 7.5, 300 mM NaCl, 50 mM imidazole pH 7.5, 0.03% (w/v) LMNG, 0.003% (w/v) CHS] at 2 mL/min each. The protein was eluted from the protein with elution buffer [25 mM HEPES pH 7.5, 300 mM NaCl, 300mM imidazole pH 7.5, 10% (v/v) glycerol, 0.003% (w/v) LMNG, 0.0003% (w/v) CHS] at a flowrate of 0.5. mL/min. The Buffer was exchanged to storage buffer [25 mM HEPES pH 7.5, 300 mM NaCl, 10% (v/v) glycerol, 0.003% (w/v) LMNG, 0.0003% (w/v) CHS] using PD-10 desalting columns and concentrated to 30 mg/mL using Amicon concentrators at 1200 g. The purified protein was flash-frozen in liquid nitrogen and stored at -80°C.

### Crystallization

In preparation for crystallization, samples of β_2_AR in storage buffer at a concentration of 30 mg/mL were thawed and supplemented with 2 mM of photoazolol-1 (PZL-1). Two parts of the protein solution were mixed with three parts of molten monoolein [supplemented with 10% (w/v) cholesterol] and homogenized until a clear phase was generated. Initial LCP screening was done using a Gryphon robot (ARI) in 96-well glass sandwich plates (Laminex). 40 nL of LCP were placed in each well and covered with 800 nL of mother liquor supplemented with 10 μm photoazolol-1. Several commercial screens were used to find initial crystallization conditions. After optimization, crystallization was scaled up using a modified in-well crystallization method^70^ in EasyXtal plates (Molecular Dynamics). This allowed reproducible production of large amounts of crystal-laden LCP needed for serial crystallography. 10 μL of LCP were added to wells containing 300 μl of mother liquor [100 mM tri-sodium citrate pH 6.2, 245 mM Li_2_SO_4_, 32% (w/v) PEG 350 MME, 10 μM photoazolol-1] at a 1:30 LCP:mother liquor ratio and the plate was shielded against ambient illumination with aluminum foil and incubated at 16°C. First crystals appeared within 24 hr and were fully grown after 96 hr. The crystal-laden LCP was harvested and pooled in 500 μL Hamilton syringes and topped off with small amounts of mother liquor to avoid drying. All crystallization steps were conducted under red-light conditions and at 20°C ambient temperature.

### Time-resolved data collection

The 17 ns time-point was collected using the coherent X-ray imaging (CXI) beamline at the Linac Coherent Light Source (LCLS). Samples were delivered to the X-ray using high viscosity extrusion^71^. The crystal-laden LCP was modified by adding PEG2000 and monoolein to reach a viscosity suitable for jetting. The final jetting phase contained 62.5% crystal-laden LCP, 34.7% monoolein and 2.8% PEG2000. Jetting was conducted with a 75 μm glass capillary nozzle at 21.5 μm/ms, corresponding to 180 μm between consecutive XFEL pulses. Data was collected using a Jungfrau 4M detector at 120 Hz with an X-ray focused to about 2×2 μm (horizontal x vertical) FWHM at 8.8keV. A pump laser diode (EKSPLA NL-204) at 355 nm was used to trigger the isomerization reaction. The laser spot size was 88 μm (1/e^2^) and delivered a fluence of 216 mJ/cm^2^ in 8 ns pulses. The light dataset was recorded with a 1:1 dark to light ratio with the dark images being goose triggered, meaning that the laser arrives on the interaction zone 40 ns after the X-ray pulse. True-dark data were recorded before the pump-probe experiment for control.

The slow time-point (∼10 s) was collected at the Cristallina experimental station of SwissFEL. Here, samples were delivered with solid support, using SOS chips^72^. Each chip was loaded with a volume of 25 μl of LCP sample and probed with 360x360 (columns x rows) XFEL pulses. Data were collected using a Jungfrau 8M detector at 100 Hz with an X-ray focused to 5x5 μm FWHM at 12.044 keV. A laser diode at 405 nm was used for illumination and set to a 225x225 μm spot size with a measured power of 100 mW. Due to the scan pattern, area of illumination and repetition rate, this measurement results in a ∼10 s time point (**Supplementary Figure 2**).

### Structure determination and refinement

The data collected was processed with CrystFEL 0.10.2^73,74^ using peakfinder8 for peak detection and XGANDALF^75^ for indexing. For Cristallina data, the optimal settings were determined to be: *--threshold=20 –min-snr=4*.*0, --min-pix-count=1*.The data was merged and scaled using partialator with the option *-m unity -n 1 --push-res=1*.*5* and the CCP4 suite ^76^ used to create MTZ files. Datasets were further treated for anisotropy with the SATARANISO server^77^. For LCLS data, data was merged and scaled using partialator with the option *-m xsphere -n 1* and the CCP4 suite^76^ was used to create MTZ files. Dark and light datasets were scaled together during the partialator step and then split using the custom-split option.

To solve the initial models, molecular replacement with the search model 2RH1 was used. To refine the models, iterative cycles of phenix.refine^78^ and manual rebuilding in coot were done. Xtrapol8^79^ was used to calculate q-weighted difference maps and q-weighted extrapolated maps with standard settings. For LCLS, fewer crystals patterns were collected than for the Cristallina experiment, therefore extrapolation was based on F*calc* of the dark model to improve the quality of the extrapolated map. After visual examination, the activation level was determined to be around 28%. For Cristallina data, the extrapolation was carried from F*obs* from the dark dataset and the activation level was manually determined to be around 22 %. The coordinates of light-activated models were refined against their respective extrapolated maps. After refinement, a mixed model was created for both experiments, containing the appropriate Dark models and Light models according to their determined occupancy. These mix-models were ultimately refined [B-factor and TLS^80^] against their respective light-activated datasets (non-extrapolated). Ligand restraints were generated using eLBOW^81^. The protein model quality was controlled with MolProbity^82^.

### Molecular Mechanics/Generalized Born Surface Area (MM/GBSA) calculations

Ligand binding energies were estimated using the MM/GBSA method. For this, both receptor ligand complexes (dark and light) were prepared using protein preparation wizard in Schrödinger Maestro^83,84^ using default settings (adding hydrogens, set charges for ionizable groups at pH 7.4 using PropKa and Epik). MM/GBSA calculations were run using Prime on the prepared complexes^84-86^, including an area of 5 Å around the ligand center allowing for protein flexibility to counteract the distorted cis ligand conformation.

### Cell Culture

The activity of photoswitchable ligands against β_2_AR was evaluated using HEK293 H188 M1 cells, which stably express a cAMP Epac-S^H188^ FRET biosensor^87^ and was reported in our previous paper^**27**^. We maintained the cell line stably expressing the Epac-SH^188^ cAMP biosensor at 37ºC, 5% (w/v) CO_2_ in 4,5 g/L D-glucose Dulbecco’s Modified Eagle Medium (DMEM, GIBCO) supplied with 10% heat inactivated FBS (GIBCO) and 1% penicillin-streptomycin (10,000 U/mL, GIBCO). All assays were performed at room temperature. Adherent cells were grown in 150-mm dishes to 75-90% confluence and recovered by rinsing once with PBS (GIBCO), followed by incubation with Trypsin-EDTA (Sigma-Aldrich) for 5 min until detachment of cells was observed. Cells were then centrifugated; in parallel, 10 µL of the single cell suspension were counted using a Neubauer Chamber. The supernatant was carefully removed, and cells were resuspended in DMEM complete medium to obtain a solution at 1.0 × 10^6^ cells/mL. 100,000 cells per well were seeded in a transparent 96-well microplate (Thermo Scientific Nunc Microwell) and left at 37ºC with 5% CO_2_ for approximately 24h. The cAMP EPAC sensor buffer (14 mM NaCl, 50 nM KCl, 10 nM MgCl_2_, 10 nM CaCl_2_, 1 mM HEPES pH 7.2, 1.82 mg/mL Glucose,) ^87^ supplemented with 100 µM IBMX was used as the assay medium in all FRET-based experiments. Fluorescence values were measured using a Tecan Spark M20 multimode microplate reader equipped with the Fluorescence Top Standard Module and defined wavelength settings (excitation filter 430/20 nm and emission filters 485/20 nm and 535/25 nm). FRET ratio was calculated as the relation of the fluorescent donor emission (td^cp173^V, 485 nm) divided by the acceptor emission (mTurq2Δ, 535 nm). The FRET ratio was normalized to the effect of the buffer (0%) and the maximum response obtained with cimaterol (100%). External light was applied using the 96-well LED array plate (LEDA Teleopto). Each set of experiments was performed three to five times with each concentration in duplicate or triplicate.

### Dose-Response Assays

To perform functional assays with azobenzene ligands in HEK293 H188 M1 cells we prepared two different plates, one for each light condition. For all assays, both plates were left to incubate with the studied compounds for 1h at room temperature. To induce photoswitching, the “light plate” was exposed to continuous illumination (380 nm) during 15 min using the LED array plate (LEDA Teleopto). Fluorescence values were thereafter measured and, subsequently, the cells were washed three times with assay buffer. Dose-response curves for each compound were obtained using a constant concentration of the agonist cimaterol (10 nM) in the dark and upon illumination.

### Binding competition experiments

The binding of photoswitchable ligands towards β_2_AR was evaluated using HEK293 cells transiently transfected with β_2_AR with lipofectamine. To perform binding competition assays a carazolol-based fluorescent ligand Carazolol-KK114 (Car-KK114) which competes with the azobenzene ligands for the same β_2_AR binding site was used^88^. To evaluate the binding affinites of the photoswitchable ligands we prepared two different plates, one for each light condition. Both plates were co-incubated with 100 nM of Car-KK114 and different concentrations of the studied compounds for 1h at room temperature. Subsequently, the cells were washed three times with assay buffer and fluorescence values were measured. To induce photoswitching, one plate was exposed to continuous illumination at 380 nm during 15 min using the LED array plate (LEDA Teleopto), while another plate was left under dark conditions for 15 min. Fluorescence values were thereafter measured and, subsequently, the cells were washed and incubated with 100 nM of Car-KK114 for 1h at room temperature. Finally, the cells were washed three times with assay buffer and fluorescence values were measured.

### Data analysis

All experiments were analyzed using GraphPad Prism 10.4.0 (GraphPad Software, San Diego, CA). Stimulation dose–response data was fitted using the log(agonist) vs response (three parameters) function. Inhibition dose–response data was fitted using the log (antagonist) vs response (three parameters) function. Competitive binding data was fitted using the One site – Fit Ki function.

### Synthesis materials and methods

All starting materials were obtained from commercial sources and used without further purification. Anhydrous solvents were obtained from a solvent purification system (*PureSolv-EN*^*TM*^) and kept under a nitrogen atmosphere. Reactions were monitored by thin layer chromatography (TLC) on silica gel (60F, 0.2 mm, ALUGRAM Sil G/UV254 *Macherey-Nagel*) and visualized with 254 nm UV light. Reactions under microwave irradiation were carried out in a *CEM Discover Focused™* Microwave reactor. This instrument is constituted by a continuous focused microwave power delivery system with selectable power output (0-300 W). Reactions were performed in 5 mL sealed glass vessels. Temperature of the vessel content was monitored using an IR sensor and the indicated temperature corresponds to the maximal temperature reached during each experiment. Reaction vessels were magnetically stirred by means of a rotating magnetic plate located below the floor of the microwave cavity. The specified time corresponds to the total irradiation time. Efficient cooling was accomplished by means of pressurized air during the entire experiment. Flash column chromatography was performed using silica gel 60 (*Panreac*, 40-63 μm mesh) or by means of RediSep Silica (*Biotage*) and/or RediSep HP C18 Gold (*Biotage*) columns, automated with Isolera One with UV-Vis detection (*Biotage*). Nuclear Magnetic Resonance (NMR) spectroscopy was performed using a *400 MHz Brüker Avance NEO 400 MHz* spectrometer. Chemical shifts are reported in δ (ppm) relative to the residual non-deuterated solvent signal (CDCl3 δ = 7.26 ppm (^1^H), δ = 77.16 ppm (^13^C); DMSO-*d6* δ = 2.50 ppm (^1^H), δ = 39.51 ppm (^13^C), CD3OD δ = 3.31 ppm (^1^H), δ = 49.3 ppm (^13^C)). The following abbreviations have been used to designate multiplicities: s=singlet, d=doublet, t=triplet, q=quadruplet, qu=quintuplet, h=heptet, m=multiplet, br=broad signal, dd=doublet of doublet, ddd=doublet of doublet of doublet, dddd=doublet of doublet of doublet of doublet, dt=doublet of triplet, qd=quadruplet of doublet. Coupling constants (*J*) are reported in Hz. High-resolution mass spectra (HRMS) and elemental composition were performed on a FIA (Flux Injected Analysis) with Ultrahigh-Performance Liquid Chromatography (UPLC) *Acquity Premier (Waters)* coupled to LCT Premier Select Series Cyclic IMS (*Waters*). Data from mass spectra was analyzed by electrospray ionization in positive mode using MassLynx 4.2 Software (*Waters*). Spectra were scanned between 50 and 1200 Da with values every 0.4 seconds and peaks are reported as *m/z*. Purity of final compounds was determined by High-Performance Liquid Chromatography (HPLC). Analytical HPLC was performed on a *Thermo Ultimate 3000SD* (*Thermo Scientific Dionex*) coupled to a PDA detector and Mass Spectrometer *LTQ XL ESI-ion trap* (*Thermo Scientific)* (HPLC-PDA-MS)) or on a *Waters 2795 Alliance* coupled to a DAD detector (*Agilent 1100*) and an *ESI Quattro Micro* MS detector (*Waters*); HPLC columns used were *ZORBAX Eclipse Plus C18* (4.6x150mm; 3.5μm) and *ZORBAX Extend-C18* (2.1 x 50 mm, 3.5 μm) respectively. HPLC purity was determined using the following binary solvent system: 5% acetonitrile (v/v) in 0.05% formic acid (v/v) for 0.5 minutes, from 5 to 100% acetonitrile (v/v) in 5 minutes, 100% acetonitrile (v/v) for 1.5 minutes, from 100 to 5% acetonitrile (v/v) in 2 minutes and 5% acetonitrile (v/v) for 2 minutes. The flow rate was 0.5 mL/min, column temperature was fixed to 35 ºC and wavelengths from 210-600 nm were registered. Purity determination was performed with Liquid Chromatography (LC) coupled to a photodiode detector (PDA) and a mass spectrometer (MS). Three different equipment with different methods have been used and are described as followed. Waters 2795 Alliance separation module coupled to a diode array detector (Agilent 1100) scanning at a wavelength range of 210-600 nm and an ESI Quattro Micro MS detector (Waters) in positive mode with mass range (m/z) of 150-1500. A column ZORBAX Extend-C18 3.5 μm 2.1x50mm (Agilent) at 35ºC was used with a mixture of A = H_2_O + 0.05% formic acid and B = MeCN + 0.05% formic acid as mobile phase and the method as follows: flow 0.5 mL/min, Gradient t = 0.0 min 5% B, t = 0.5 min 5% B, t = 5.5 min 100% B, t = 7.0 min 100% B, t = 8.0 min 5% B, t = 10.0 min 5% B, total runtime: 10 min.

### Synthesis of fluorescent ligand Car-KK114

For the binding experiments the fluorescent ligand Car-KK114 previously reported by Mitronova et al. was used^88^. In addition, Car-KK114 was synthesized following the conditions depicted in Scheme 1. 4-hydroxycarbazol (**1**) was alkylated with glycidyl tosylate (**2**) to form 4-(glycidyloxy)carbazole (**3**). Next, the nucleophilic epoxide ring opening with *tert*-butyl N-[3-amino-3-methylbutyl] carbamate (**4**) followed by the amine deprotection gave the carazolol derivative **6**. Subsequently, a PEG linker was introduced followed through an amide formation with the azido activated carboxylic acid **7** and the resulting azide **8** was reduced by hydrogenation to give the carazolol derivative **9** with a terminal primary amino group, which was used to attach of the fluorescent dye KK114 and form the final Car-KK114 (**10**).

**Scheme 1:**
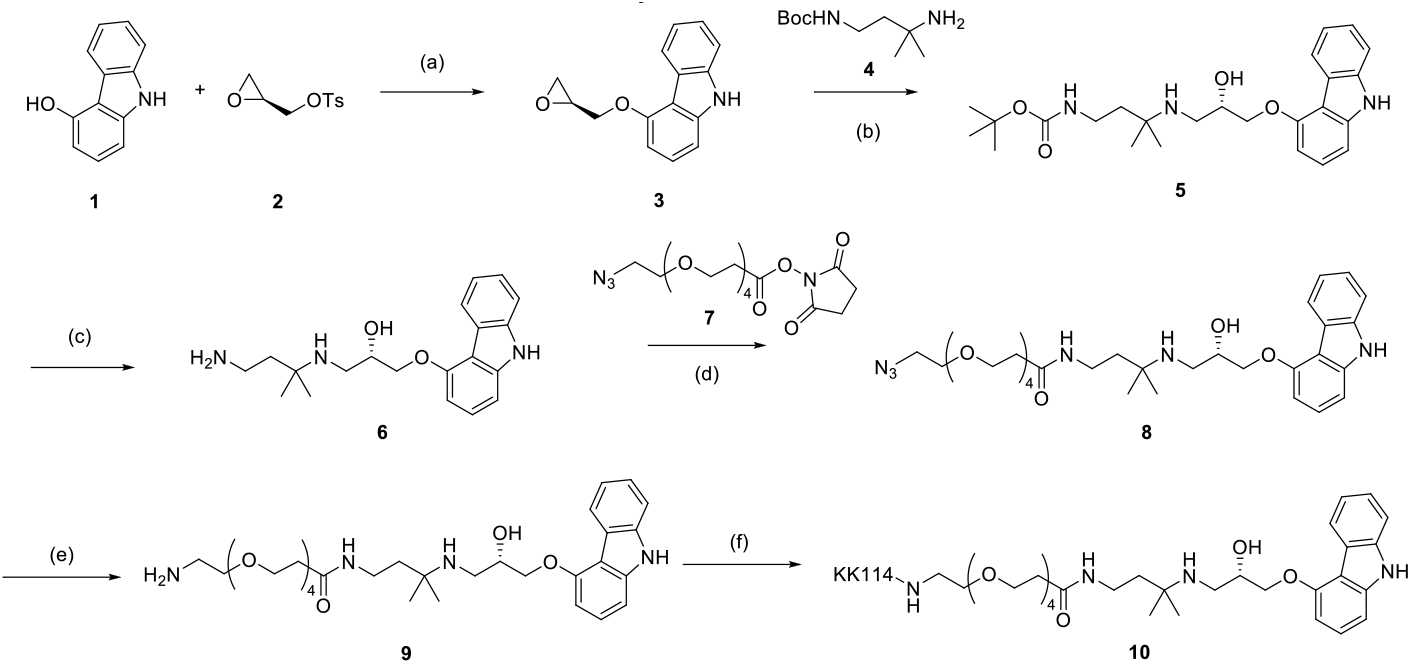
Synthesis of fluorescent ligand Car-KK114. (a) DMF, rt, ON, 48%; (b) iPrOH, 100 ºC, μW 2h, 76%; (c) TFA, DCM, rt, 6h, 56%; (d) TEA, DMF, 60 ºC, 16h, 56%; (e) H_2_, Pd/C, HCl, MeOH, 3 bar, 3h, 16%.; (f) KK114-NHS, TEA, DMSO, rt, 3h, 33%.

#### *(S)*-4-(oxiran-2-ylmethoxy)-9H-carbazole (3)

9H-carbazol-4-ol (**1**, 500 mg, 2.73 mmol) and CS_2_CO_3_ (1.33 g, 4.1 mmol) were dissolved in dry DMF (3 ml) and the reaction mixture was stirred for 10 minutes. *(S)*-oxiran-2-ylmethyl 4-methylbenzenesulfonate (**2**, 934 mg, 4.1 mmol) was added and the reaction mixture was stirred at room temperature overnight. Afterwards, the reaction mixture was concentrated under reduced pressure. Saturated aqueous solution of NaHCO_3_ and EtOAc were added and the layers were separated. The aqueous layer was extracted with EtOAc twice. The combined organic layers were washed with brine and dried over MgSO_4_. The crude was purified by automated normal phase *flash* column chromatography (from Hex:EtOAc 85:15 to 25% EtOAc) to give the title compound as a brown solid (653 mg, 48%). <u>^1^H NMR</u> (400 MHz, CDCl_3_) δ 8.34 (d, *J* = 7.8 Hz, 1H), 8.07 (s, 1H), 7.43 – 7.38 (m, 2H), 7.32 (t, *J* = 8.0 Hz, 1H), 7.06 (d, *J* = 8.1 Hz, 1H), 6.66 (d, *J* = 8.0 Hz, 1H), 4.47 (dd, *J* = 11.0, 3.3 Hz, 1H), 4.27 (dd, *J* = 11.0, 5.4 Hz, 1H), 3.60 – 3.52 (m, 1H), 3.00 (t, *J* = 4.5 Hz, 1H), 2.90 (dd, *J* = 4.9, 2.6 Hz, 1H). ^1^H NMR signals match those reported in literature^89^. HPLC-PDA-MS: RT = 3.64 min, λ_max_ = 242 nm, [M+H]^+^ = 240.32; purity (254 nm): 100%.

#### *tert*-butyl (3-amino-3-methylbutyl)carbamate (4)

di-*tert*-butyl dicarbonate (1.18 g, 5.4 mmol) was dissolved in DCM (5 ml). To the ice-cooled mixture, 3-methylbutane-1,3-diamine (1.18 ml, 11.34 mmol) and TEA (1.58 ml, 11.34 mmol) were added and the reaction mixture was stirred at room temperature overnight. Saturated aqueous solution of Na_2_CO_3_ and DCM were added and the layers were separated. The organic layer was washed with saturated aqueous solution of Na_2_CO_3_ twice. The combined organic layers were washed with brine and dried over MgSO_4_. The crude was purified by automated normal phase *flash* column chromatography (MeOH:DCM 90:10%) to give the title compound as a white solid (1 g, 98%). <u>^1^H NMR</u> (400 MHz, CDCl_3_) δ 4.93 (s, 1H), 3.01 (d, J = 6.3 Hz, 2H), 1.45 (s, 9H), 1.10 (s, 6H). ^>1^>H NMR signals match those reported in literature^90^.

#### *tert*-butyl *(S)*-(3-((3-((9H-carbazol-4-yl)oxy)-2-hydroxypropyl)amino)-3-methylbutyl)carbamate (5)

4-(oxiran-2-ylmethoxy)-9H-carbazole (**3**, 300 mg, 1.25 mmol) was dissolved in iPrOH (2 ml). *tert*-butyl (3-amino-3-methylbutyl)carbamate (330 mg, 1.63 mmol) was added and the reaction mixture was stirred at 100 ºC for 2 h at the microwaves. Afterwards, the reaction mixture was concentrated under reduced pressure and the crude was purified by automated reverse-phase *flash* column chromatography (from H_2_O:ACN 95:5 + 0.1% HCOOH to 100% ACN + 0.1% HCOOH to give the title compound as a white solid (420 mg, 76%). <u>^1^H NMR</u> (400 MHz, CDCl_3_) δ 8.26 (d, *J* = 7.8 Hz, 1H), 8.23 (s, 1H), 7.43 – 7.35 (m, 2H), 7.32 (t, *J* = 8.0 Hz, 1H), 7.22 (ddd, *J* = 8.1, 6.4, 1.7 Hz, 1H), 7.06 (d, *J* = 8.0 Hz, 1H), 6.67 (d, *J* = 8.0 Hz, 1H), 5.27 (s, 1H), 4.34 – 4.27 (m, 1H), 4.27 – 4.19 (m, 2H), 3.26 – 3.17 (m, 2H), 3.04 – 2.97 (m, 1H), 2.92 – 2.85 (m, 1H), 1.64 – 1.56 (m, 2H), 1.43 (s, 9H), 1.14 (s, 6H). <u>^13^C NMR</u> (101 MHz, CDCl_3_) δ 155.20, 141.10, 138.89, 126.83, 125.16, 123.01, 122.61, 119.78, 112.83, 110.24, 104.05, 101.39, 70.45, 69.34, 52.75, 44.82, 39.96, 36.96, 28.59, 27.17. HPLC-PDA-MS: RT = 3.24 min, λ_max_ = 242 nm, [M+H]^+^ = 442.48; purity (254 nm): 100%.

#### *(S)*-1-((9H-carbazol-4-yl)oxy)-3-((4-amino-2-methylbutan-2-yl)amino)propan-2-ol (6)

Trifluoroacetic acid (2.71 ml, 35.2 mmol) was added dropwise to an ice-cooled solution of compound **5** (420 mg, 0.95 mmol) in DCM (2 ml). The reaction mixture was stirred at room temperature for 6 h and was concentrated under reduced pressure to give a crude that was purified by automated reverse-phase *flash* column chromatography (from H_2_O:ACN 95:5 + 0.1% HCOOH to 100% ACN + 0.1% HCOOH to give the title compound as a white solid (181 mg, 56%). <u>^1^H NMR</u> (400 MHz, MeOD) δ 8.26 (d, *J* = 7.8 Hz, 1H), 7.43 (d, *J* = 8.1 Hz, 1H), 7.37 – 7.32 (m, 1H), 7.30 (t, *J* = 7.5 Hz, 1H), 7.18 – 7.12 (m, 1H), 7.10 (d, *J* = 8.1 Hz, 1H), 6.71 (d, *J* = 7.9 Hz, 1H), 4.50 – 4.43 (m, 1H), 4.40 – 4.34 (m, 1H), 4.30 – 4.24 (m, 1H), 3.53 – 3.46 (m, 1H), 3.35 – 3.26 (m, 1H), 3.12 – 3.03 (m, 2H), 2.15 – 2.07 (m, 2H), 1.44 (s, 3H), 1.43 (s, 3H). <u>^13^C NMR</u> (101 MHz, MeOD) δ 155.98, 142.96, 140.76, 127.45, 125.81, 123.64, 123.30, 119.82, 113.50, 111.31, 105.46, 101.65, 70.75, 67.27, 59.51, 45.88, 36.17, 35.99, 23.14, 23.13. HPLC-PDA-MS: RT = 2.31 min, λ_max_ = 242 nm, [M+H]^+^ = 342.46; purity (254 nm): 100%.

#### *(S)*-N-(3-((3-((9H-carbazol-4-yl)oxy)-2-hydroxypropyl)amino)-3-methylbutyl)-1-azido-3,6,9,12-tetraoxapentadecan-15-amide (8)

The amine **6** (181 mg, 0.53 mmol) and N_3_-PEG_4_-(CH_2_)_2_CO-NHS ester (**7**, 206 mg, 0.53 mmol) were dissolved in dry DMF (2 ml), anhydrous TEA was added and the reaction mixture was stirred at 60 ºC overnight. Afterwards, the reaction mixture was concentrated under high vacuum and purified by automated normal phase *flash* column chromatography (from DCM:MeOH 90:10 to 100% DCM) to give the title compound as a white solid (181 mg, 56%). <u>^1^H NMR</u> (400 MHz, CDCl_3_) δ 9.09 (s, 1H), 8.25 (d, *J* = 7.8 Hz, 1H), 7.41 – 7.29 (m, 2H), 7.29 – 7.23 (m, 1H), 7.21 – 7.14 (m, 1H), 7.02 (d, *J* = 8.1 Hz, 1H), 6.60 (d, *J* = 8.0 Hz, 1H), 4.27 – 4.21 (m, 1H), 4.21 – 4.13 (m, 2H), 3.64 – 3.49 (m, 16H), 3.29 – 3.25 (m, 2H), 3.25 – 3.19 (m, 2H), 2.98 – 2.93 (m, 1H), 2.87 – 2.80 (m, 1H), 2.30 (t, *J* = 6.0 Hz, 2H), 1.54 – 1.47 (m, 2H), 1.07 (s, 6H). <u>^13^C NMR</u> (101 MHz, CDCl_3_) δ 171.37, 155.06, 141.20, 139.01, 126.60, 124.90, 122.81, 122.33, 119.34, 112.47, 110.37, 104.12, 100.93, 70.57, 70.54, 70.49, 70.38, 70.28, 70.21, 70.07, 69.90, 69.39, 67.33, 52.44, 50.57, 44.81, 38.89, 37.02, 35.77, 27.15, 27.09. HPLC-PDA-MS: RT = 2.91 min, λ_max_ = 242 nm, [M+H]^+^ = 615.60; purity (254 nm): 98.5%.

#### *(S)*-N-(3-((3-((9H-carbazol-4-yl)oxy)-2-hydroxypropyl)amino)-3-methylbutyl)-1-amino-3,6,9,12-tetraoxapentadecan-15-amide (9)

A mixure of azido compound **8** (220 mg, 0.36 mmol), Pd/C (19 mg, 10 % Wt, 0.02 mmol) and HCl (91 µl, 37% in H_2_O, 1.11 mmol) were suspended in MeOH (4 ml) and a hydrogen atmosphere of 3 bar was applied for 3 h at room temperature. The reaction mixture was filtered through Celite, washed with MeOH and concentrated under reduced pressure to give a crude that was purified by automated reverse-phase *flash* column chromatography (from H_2_O:ACN 95:5 + 0.1% HCOOH to 100% ACN + 0.1% HCOOH to give the title compound as a white solid (35 mg, 16%). <u>^1^H NMR</u> (400 MHz, MeOD) δ 8.28 (d, *J* = 7.7 Hz, 1H), 7.46 (d, *J* = 8.1 Hz, 1H), 7.36 (t, *J* = 7.1 Hz, 2H), 7.32 (t, *J* = 8.0 Hz, 1H), 7.17 (t, *J* = 7.4 Hz, 1H), 7.12 (d, *J* = 8.1 Hz, 1H), 6.74 (d, *J* = 7.9 Hz, 1H), 4.56 – 4.47 (m, 1H), 4.44 – 4.37 (m, 1H), 4.34 – 4.27 (m, 1H), 3.73 – 3.65 (m, 4H), 3.64 – 3.49 (m, 14H), 3.38 – 3.33 (m, 2H), 3.09 (t, *J* = 4.9 Hz, 2H), 2.46 (t, *J* = 5.9 Hz, 2H), 2.00 – 1.92 (m, 2H), 1.47 (s, 3H), 1.46 (s, 3H). <u>^13^C NMR</u> (101 MHz, MeOD) δ 174.40, 156.00, 142.95, 140.76, 127.50, 125.82, 123.70, 123.32, 119.86, 113.49, 111.36, 105.46, 101.68, 71.29, 71.22, 71.16, 71.12, 70.77, 70.75, 68.09, 67.69, 67.38, 60.11, 45.85, 40.46, 38.25, 37.38, 35.64, 23.79, 23.76. HPLC-PDA-MS: RT = 2.54 min, λ_max_ = 242 nm, [M+H]^+^ = 589.49; purity (254 nm): 100 %.

### Carazolol-KK114 (10)

The amine **9** (4.81 mg, 7.69 µmol), a solution of KK114-NHS ester 10,1 mM in DMSO (700 µl, 7.69 µmol) and TEA (23 µl, 11.54 µmol) were dissolved in DMSO (1.0 ml) and the reaction mixture was stirred at room temperature for 3 h. Afterwards, the crude was purified by automated reverse-phase *flash* column chromatography (from H_2_O:ACN 85:15 0.1% HCOOH to 45% ACN + 0.1% HCOOH to give the title compound as a blue solid (3.3 mg, 30%). HPLC-PDA-MS: RT = 3.10 min, λ_max_ = 222 nm, [M+H]^+^ = 1459.82; purity (254 nm): 100%. HRMS (m/z): [M+H]^2+^ calcd for C_73_H_89_F_4_N_7_O_16_S2^2+^ 729.7866, found 729.7886.

## Acknowledgments

We are grateful for the support from the PSI Crystallization Facility and the Macromolecular Crystallography group during the growing and testing of crystals at the Swiss Light Source. The Biomolecular Structure and Mechanism Program of the Life Science Zürich Graduate School is acknowledged for its academic framework for our graduate students. Use of the Linac Coherent Light Source (LCLS), SLAC National Accelerator Laboratory, is supported by the U.S. Department of Energy, Office of Science, Office of Basic Energy Sciences under Contract No. DE-AC02-76SF00515. This work was supported by the National Institutes of Health grant S10 OD025079. Beamtime at the Cristallina station of the Swiss X-ray Free Electron laser (SwissFEL) was granted under proposal 20231114 in the first user run.

## Funding

Swiss National Science Foundation Project Grants 310030_197674 (to T.W.) and 310030_207462 (J.S.)

Swiss Innovation Agency Innosuisse Grant 42711.1 IP-LS (to M.H. and J.S.)

Swiss National Science Foundation Sinergia Grant CRSII5_213507 (J.S.)

Swiss Nanoscience Institute SNI #1904 (M.C.)

PSI Research Grant (to J.B. and J.S.)

Ministerio de Ciencia e Innovación and Agencia Estatal de Investigación and ERDF - A way of making Europe, PID2020-120499RB-I00 (AL and XR) and RYC2020-029485-I (XR)

Ministerio de Ciencia, Innovación y Universidades, PID2023-152950OB-I00 (AL) and PID2023-151614NB-I00 (XR)

Agència de Gestió d’Ajuts Universitaris i de Recerca (AGAUR) - Generalitat de Catalunya, 2021 SGR 00508 (AL)

The Spanish National Research Council, 20228AT014 (XR)

## Author contributions

Conceptualization: MH, AI, XR and JS

Methodology: RS, QB, MT, RSI, XC, GG, PS, ADC, MTI, CS, TW, SM and JB

Investigation: RS, QB, MT, ADC, MTI, HG, FS, HPS, MW, JC, MC, SH, MM, TM, YK, MA, TW, RC, CS, SM, JB, MK, XR and JS

Visualization: RS, QB, MH, XR and JS

Supervision: JB, SM, MK, MH, XR and JS

Writing - original draft: RS, XR and JS

Writing - review & editing: all authors

## Data and materials availability

Coordinates and structure factors have been deposited in the PDB database under accession codes 9RKF for the Dark state recorded at SwissFEL, 9RKG for the Light activated state recorded 10 s after activation at SwissFEL, 9RKH for the Dark state recorded at LCLS, and Finally, 9RKI for the Light activated state recorded 17 ns after activation at LCLS.

