## Supplementary Figures for "Time-resolved structures of β_2_-adrenergic receptor modulation by a photoswitchable beta-blocker"

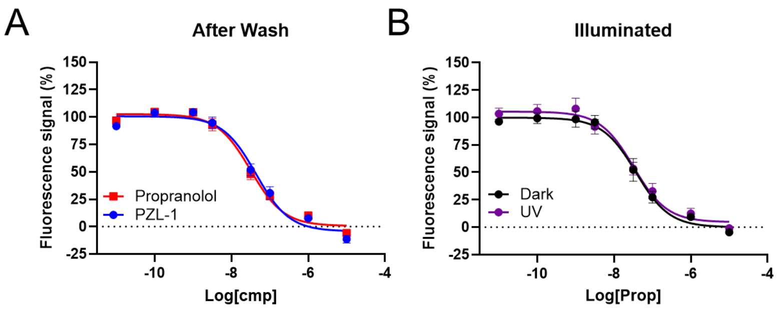


**Supplementary Figure 1: Ligand binding to the $\beta_2$AR. (A)** Competitive binding curves of photoazolol-1 and propranolol with a constant concentration of the fluorescent ligand carazolol-KK114 (100 nM). Measurements were conducted right after the samples had been thoroughly washed. **(B)** Competitive binding curves of propranolol with a constant concentration of the fluorescent ligand carazolol-KK114 (100 nM). Measurements were performed 1 hour after the thorough washing. Data are shown as the mean ± standard error of the mean (SEM) of three independent experiments in duplicate.

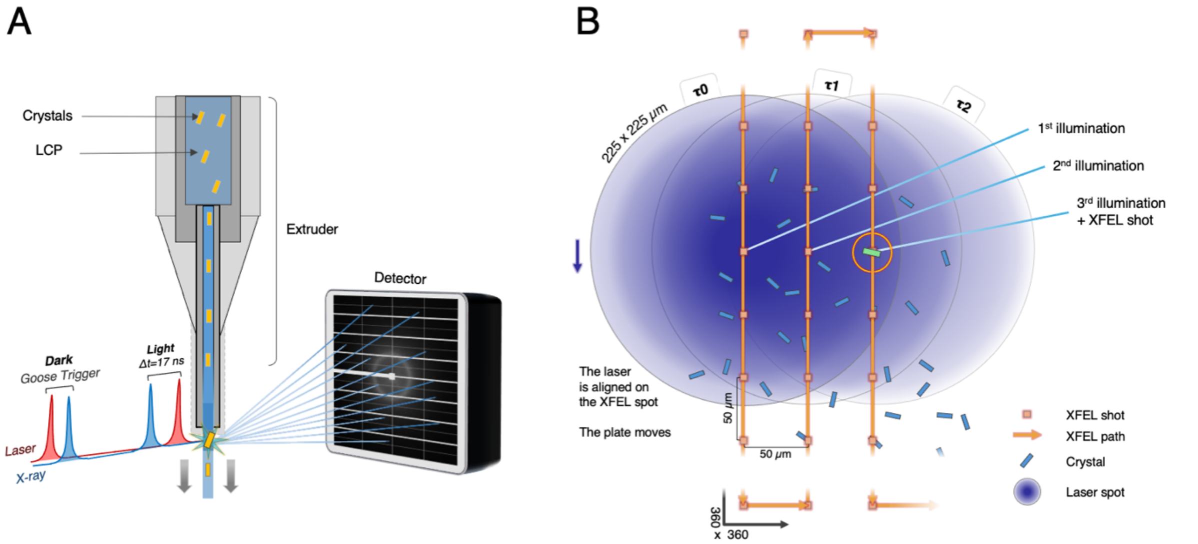


**Supplementary Figure 2: Experimental setups used for time-resolved data collection**. (**A**) The 17 ns data was collected at the Coherent X-ray Imaging (CXI) beamline at the LCLS. The classical TR-SFX setup relies on a high-viscosity extruder (HVE), similar to what we had used there previously[36,91], but relying on a nanosecond laser. The light data were recorded by triggering a nanosecond laser 17ns prior the X-ray pulse, while the dark data were collected with a "duck and goose" approach where the laser was triggered after the X-ray pulse. (**B**) The data from SwissFEL were collected at the new Cristallina fixed-target station using sheet-on-sheet devices[92]. Each chip was probed by the XFEL pulses in a 360 x 360 matrix (columns x rows), with each column taking 3.6s to complete at the 100Hz repetition rate of SwissFEL. During data collection, the sample was illuminated by a laser diode with a spot size of ≅ 225 µm diameter. Considering 50 µm distance between XFEL shots, at a time **τ**0, the laser shines up to the column $c_{+2}$ where is our crystal of interest (green crystal in the figure). Based on this experimental setup, the time between the first illumination and the X-ray probe is thus approximately 10 s. The main advantages of the solid-support system are a sample efficient data collection and the ability to probe longer time delays than are possible with extruder-based setups. The drawbacks are lower time accuracy and higher chance to measure a mixture of intermediates compared to the accurate pump-probe setup at LCLS.

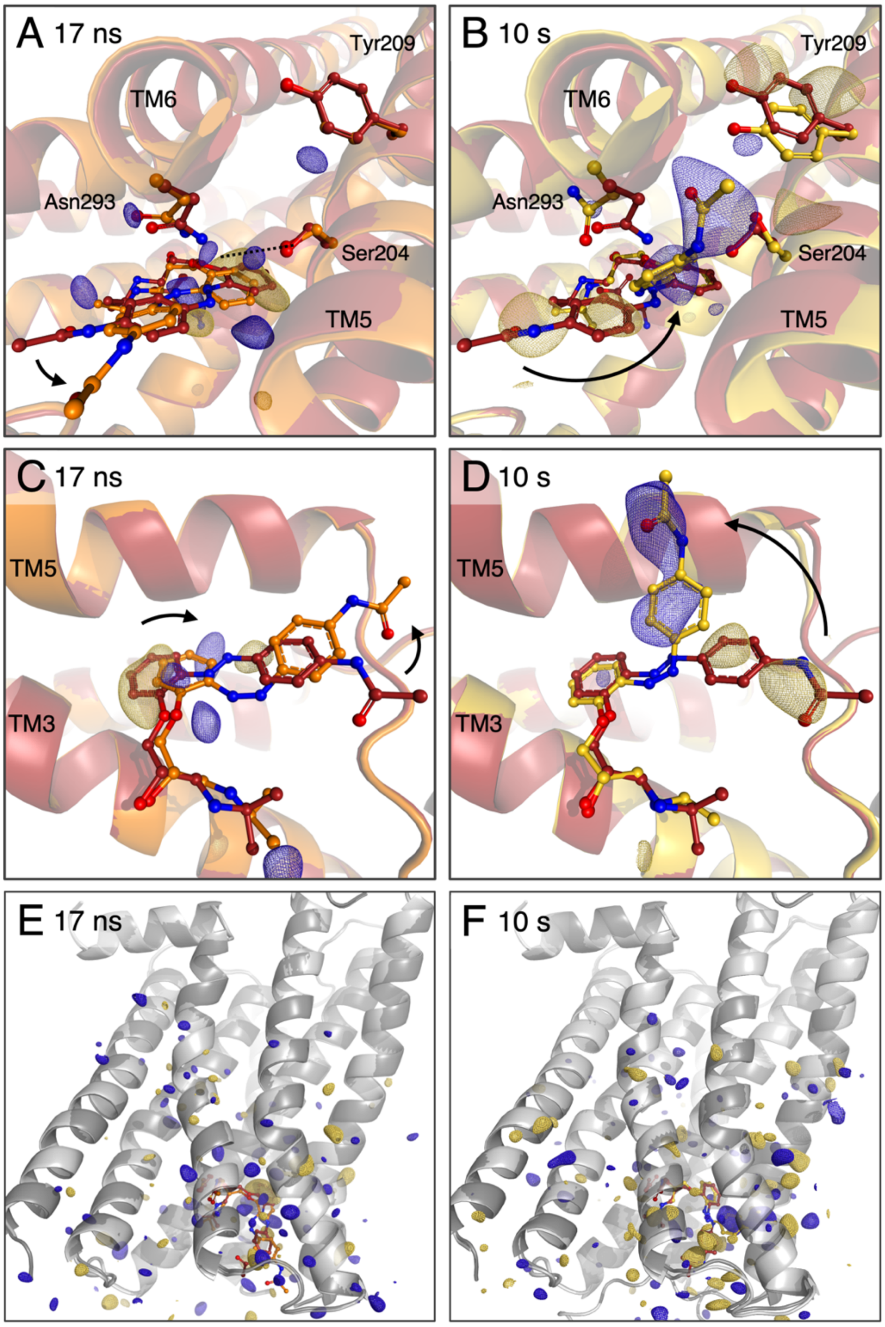


**Supplementary Figure 3: Difference electron density maps.** (**A and C**) Early rearrangements in response to photoazolol-1 isomerization. The $F_o$(17ns) − $F_o$(dark) electron density map indicates isomerization and a shift in the position of photoazolol-1 shortly after photoactivation. This transient *cis* configuration induces only a little movement in the binding pocket. (**B and D**) Adaptation of the binding pocket upon photoazolol-1 relaxation. The $F_o$(~10s) − $F_o$(dark) electron density map indicates relaxation of the initially strained *cis* configuration, including a repositioning of the p-acetamido substituted ring II of photoazolol-1 (arrows). The new position in between TM5 and TM6 interrupts the interaction between Asn293$^{6.55}$ and Ser204$^{5.43}$. **(E and F)** $F_o$(light) − $F_o$(dark) electron density maps carved at 3 angstroms around the entirety of $\beta_{2A}$R models for respectively 17 ns and 10 s. The majority of the signal is visible around the ligand binding pocket, and the stronger peaks are located on the ligand. All maps are represented at 3 sigma, with positive density in blue and negative density in yellow.

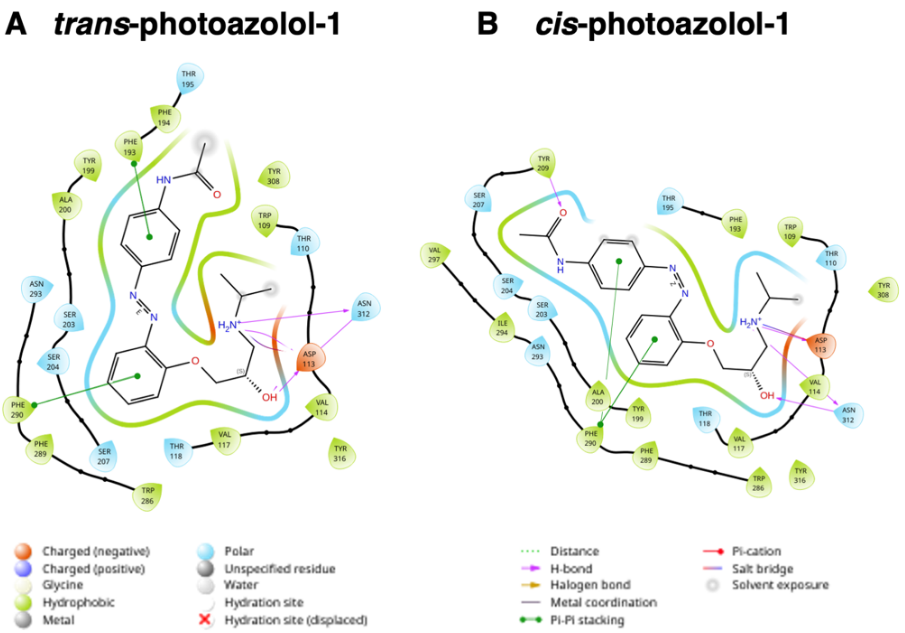


| C Ligand Binding Energy | | |
|---|---|---|
| | *trans*-photoazolol-1 | *cis*-photoazolol-1 |
| Coulomb | -62.02 | -63.16 |
| Covalent binding | -2.43 | 2.36 |
| Van der Waals | -47.05 | -48.12 |
| Lipohilic | -28.94 | -26.21 |
| G.B.E.S. | 69.05 | 67.41 |
| H-bond correction | -3.25 | -3.73 |
| $\pi$-stacking correction | -2.63 | -2.22 |
| Total energy | -77.27 | -73.68 |

**Supplementary Figure 4: Ligand binding energy and interaction patterns.** 2D projection of the interaction patterns with photoazolol-1 in it's (**A**) *trans* and (**B**) relaxed *cis* configurations. The type of interaction and the participating protein residues are color-coded according to the shown legend. Lines indicate important interactions. Presentations were generated using SiteMap in Maestro[84]. (**C**) Ligand binding energies calculated using MM/GPSA (Molecular Mechanics/Generalized Born Surface Area) in Maestro[84]. The relative energies of photoazolol-1 in the dark (left) and 10s structures (right) compared to the relaxed ligand in solvent are shown. In both cases, ligand-receptor interactions remain highly favorable, with the **cis-isoform** exhibiting a slightly lower total energy difference. Loss of **polar interactions** with Asn293$^{6.55}$ and Ser204$^{5.43}$ is compensated by **stronger interactions** with TM5, notably a **hydrogen bond with** Tyr209$^{5.48}$. Ligand **intercalation** between helices 5 and 6 enhances **van der Waals interactions**, bringing **ring II** into close contact with both helices. The hydrophobic acetate group also shifts into a **hydrophobic receptor region**, reducing unfavorable G.B.E.S. (Generalized Born electrostatic solvation) energy and increasing the contact area. A **loss of lipophilic interactions** partially offsets these favorable changes post-***trans-cis*** **isomerization**, due to helix reorganization, increasing polarity near the lipophilic isopropyl residue in the molecular fingerprint. In both cases, the **ligand-receptor complex remains in a significantly lower energy state** than the unbound ligand and receptor, aligning with the observed ligand retention in the binding pocket **several seconds after isomerization**.

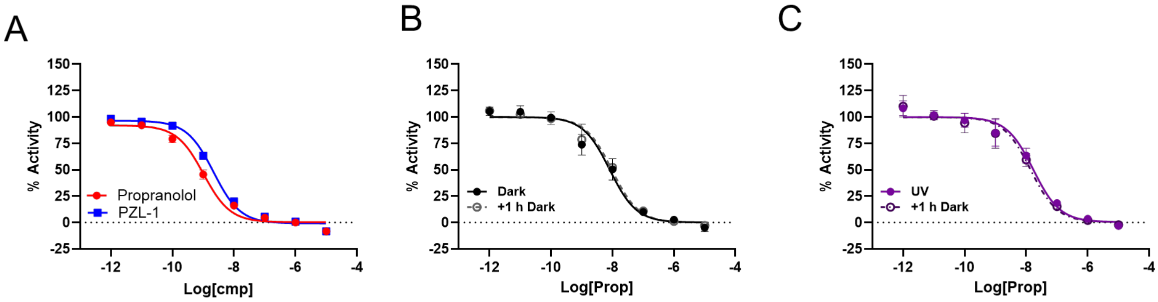


**Supplementary Figure 5: Ligand activity on β2AR before washing and after washing in the presence of Low concentration of agonist.** All experiments were performed with a constant concentration of the agonist cimaterol (10 nM). **(A)** Dose-response curves of PZL-1 and propranolol before the samples had been thoroughly washed. **(B)** Dose-response curves of propranolol 15 min after washing kept in the dark (solid black lines), and after an additional 1-hour incubation in the dark (dashed black lines). **(C)** Dose-response curves of propranolol 15 min after washing kept under light at 380 nm (solid violet lines) and after an additional 1-hour incubation in the dark (dashed violet lines). Data are shown as the mean ± SEM of three independent experiments in duplicate.

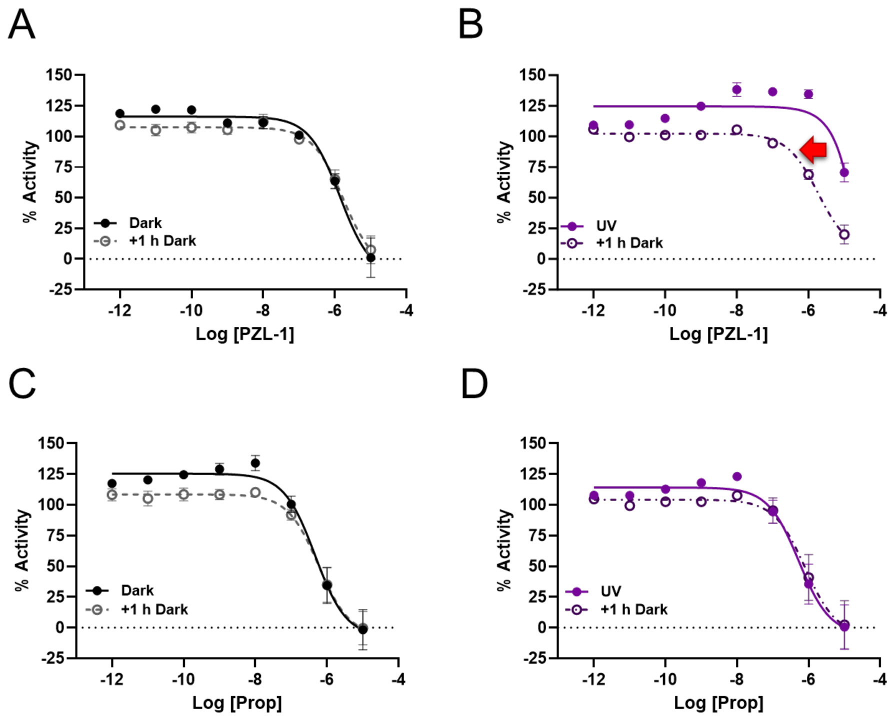


**Supplementary Figure 6: Ligands function on β2AR in the presence of high concentrations of agonist.** All experiments were performed with a constant concentration of the agonist cimaterol (1 µM). **(A)** Dose-response curves of photoazolol-1 15 min after washing kept in the dark (solid black lines), and after an additional 1-hour incubation in the dark (dashed black lines). **(B)** Dose-response curves of photoazolol 15 min after washing kept under light at 380 nm (solid violet lines) and after an additional 1-hour incubation in the dark (dashed violet lines). **(C)** Dose-response curves of propranolol 15 min after washing kept in the dark (solid black lines), and after an additional 1-hour incubation in the dark (dashed black lines). **(D)** Dose-response curves of propranolol 15 min after washing kept under light at 380 nm (solid violet lines) and after an additional 1-hour incubation in the dark (dashed violet lines). Data are shown as the mean ± SEM of three independent experiments in duplicate.

**Supplementary Movie 1: Morph between β$_2$AR structures containing dark-adapted and light-activated photoazolol-1.** This morph between the dark (red), 17 ns (orange), and 10 s (gold) structures shows how the β$_2$AR binding pocket adapts during the *trans*-to-*cis* isomerization of photoazolol-1, viewed from the extracellular side (left) and the membrane plane (right). For comparison, the dark-state structure is shown in transparent grey throughout the movie.

| Dataset | *SwissFEL – Cristallina – 10s* | | *LCLS – CXI – 17ns* | |
|---|---|---|---|---|
| | **Dark** | **Light** | **Dark** | **Light** |
| **Data Collection** | | | | |
| Space group | *C121* | | | |
| *Unit cell* *a, b, c* (Å) / α , β , γ (°) | 111.94, 172.23, 41.23 90.0, 106.2, 90.0 | | | |
| *Indexed crystals* | 88 849 | 84 524 | 55 816 | 40 650 |
| **Overall Statistics (High-Resolution Statistics)** | | | | |
| Resolution (Å) | 53.75 – 2.45 (2.54 – 2.45) | 53.75 – 2.45 (2.54 – 2.45) | 28.33 – 2.5 (2.59 – 2.5) | 28.33 – 2.6 (2.69 – 2.6) |
| Total Reflections | 13 464 666 | 12 834 526 | 3 979 447 | 2 804 141 |
| Unique Reflections | 27 481 (2741) | 27 481 (2741) | 25 868 (2609) | 22 987 (2304) |
| *<I/s(I)>* | 8.40 (0.75) | 8.27 (0.74) | 6.43 (0.54) | 6.16 (0.63) |
| $CC^*$ | 0.998 (0.626) | 0.998 (0.669) | 0.996 (0.767) | 0.995 (0.711) |
| $R_{split}$ (%) | 8.16 (143.62) | 8.35 (142.77) | 12.35 (153.68) | 13.80 (133.78) |
| Completeness (%) | 100 (100) | 100 (100) | 100 (100) | 100 (100) |
| Multiplicity | 496 (125) | 467 (125) | 153 (104) | 121 (80) |
| **Refinement** | | | | |
| Resolution Range | 45.60 – 2.45 | 45.60 – 2.45 | 28.33 – 2.5 | 28.33 – 2.60 |
| No. Reflections | 23 701 | 23 487 | 25 813 | 22 971 |
| $R_{work}$ / $R_{free}$ (%) | 18.09 / 21.86 | 18.31 / 21.69 | 19.31 / 21.83 | 19.59 / 23.40 |
| **No. Atoms** | | | | |
| Protein | 3556 | 7107 | 3560 | 7094 |
| Ligand | 342 | 696 | 323 | 667 |
| Solvent | 31 | 4 | 26 | 33 |
| **B Factors** | | | | |
| Protein | 76.74 | 78.34 | 79.54 | 93.17 |
| Ligand | 95.68 | 100.63 | 98.41 | 109.33 |
| Solvent | 67.35 | 52.42 | 72.19 | 37.97 |
| **R.m.s Deviations** | | | | |
| Bond Lengths (Å) | 0.009 | 0.004 | 0.005 | 0.004 |
| Bond Angles (°) | 0.690 | 0.782 | 0.898 | 0.853 |
| **Ramachandran** | | | | |
| Favoured (%) | 97.28 | 97.28 | 97.73 | 97.73 |
| Allowed (%) | 2.72 | 2.72 | 2.27 | 2.29 |
| Outliers (%) | 0.00 | 0.00 | 0.00 | 0.00 |
| ***PDB*** | **9RKF** | **9RKG** | **9RKH** | **9RKI** |

**Supplementary Table 1: Crystallographic data and refinement statistics.**
